# ‘Stray Appetites’: A Socio-Ecological Analysis of Free-Ranging Dogs Living Alongside Human Communities in Bangalore, India

**DOI:** 10.1101/2020.07.19.210617

**Authors:** Shireen Jagriti Bhalla, Roy Kemmers, Ana Vasques, Abi Tamim Vanak

**Author notes:** Equal contributors (Roy Kemmers, Ana Vasques, Abi Tamim Vanak). Author contributions All authors contributed to the study conception and design. Material preparation, data collection and analysis were performed by Shireen J Bhalla and supervised by Roy Kemmers, Abi T Vanak and Ana Vasques. The first draft of the manuscript was written by Shireen J Bhalla and all authors commented on previous versions of the manuscript. All authors read and approved the final manuscript.

## Abstract

Across the developing world, humans and free-ranging domestic dogs share common spaces. The relationship between these dogs and humans can range from one of dependence, to apathy, to conflict. Given the high number of humans attacked by dogs every year in India, and the lack of an effective population control strategy, we seek to provide insights into the conflict and propose alternative population management options based on reducing the carrying capacity of the environment. We used a mixed methods approach to understand both ecological and sociological underpinnings of free-ranging dog-human relationships in Bangalore, India. We conducted a photographic capture-recapture survey of free-ranging dogs to estimate population size and linked it to the availability of potential food sources. We also conducted a qualitative survey to assess attitudes of residents towards the dog population. We found that dog population varied from 192 to 1888 per square kilometre across a gradient of housing densities. The density of houses, bakeries and garbage piles were significant predictors of dog population size. Crucially, as low as 10 to 18.3% of houses supported the large population of dogs, highlighting the need for residents to act responsibly towards the dogs. Further, we found that garbage, although significant, is a secondary food source to household-maintained dogs. Since on the whole, respondents expressed the desire for a reduction in dog population, we suggest decreasing the carrying capacity of the environment by targeting these three food sources.

## Introduction

Free-ranging dogs are an integral component of the urban ecology of Indian cities. Current estimates put the total Indian free-ranging dog population at roughly 59 million (Gompper 2014). The term ‘stray dog’ in this paper is used synonymously with free-ranging dog (FRD) which is defined by Beck (1973) as “Any dog observed without human supervision on public property or on private property with immediate unrestrained access to public property” (p. 3). These dogs play several roles in the urban environment. In developing countries, Reese (2005) found that they keep trash and vermin levels down and provide companionship to people. However, they are also prone to aggression, are noisy, and are carriers of diseases - predominantly rabies. In order to reduce the risk of diseases and attacks, there is a need for research into the ecology of free-ranging dogs, which, in turn, is essential for shaping policy on the management of their populations.

About 20 million people in India are estimated to be bitten by animals every year, 91% of which are dogs, and 60-63.6% of those are free-ranging (Sudarshan et al. 2006). This amounts to roughly 10 million free-ranging dog bites a year. Hampson et al. (2015) estimated an annual incidence of 20,847 human deaths to rabies in India, which is roughly 35% of all global deaths due to rabies. This is a decrease from the 30,000 annual deaths reported for the period of 1990-2002. Sudarshan et al. (2007) attributed this to an absolute increase in the socio-economic status of the Indian population, greater awareness and better access to treatment. However, other reports have noted that there has been a steady increase in the number of dog bite cases (KIMS 2007; Menezes 2008). There is a generally acknowledged paucity of data on the subject, and neglect in the management of the disease, indicating a lack of investment by the government in the matter. The disease has been neglected partly because deaths are scattered - three-quarters of rabies deaths occur in rural areas – and it is not seen as a health crisis, unlike other epidemics (Chatterjee 2009). Another reason is because there is a lack of coordination and a comprehensive national strategy to tackle the issue, unlike in Sri Lanka and Thailand which have made more significant progress in battling rabies (Sudarshan et al. 2007). Recently, however, the Indian media has been picking up coverage of dog-related incidents, and public pressure for more effective control of dogs is mounting.

In terms of population control, national law since 2001 stipulates that free-ranging dogs can only be controlled through the Animal Birth Control policy, also called the ABC program (Animal Welfare Board of India 2001). This task is delegated to local authorities and involves capturing the dogs, taking them to a pound where they are sterilised and immunised, and returning them to the same locality where they were found. There has been little systematic verification of the success of the ABC program, but there are doubts as to its feasibility because of the high number of dogs that would need to be sterilised for there to be any sizable effect on the population (Totton et al. 2010). Coleman and Dye (1996) published a theoretical model stating that at least 70% of the total dog population would have to be vaccinated in a short period of time to eliminate or prevent rabies on at least 96.5% of occasions. This was echoed in research conducted around the ABC program in Jodhpur, by Totton et al. (2010) who performed population and demographic studies in 2005 and 2007, before and after the implementation of the ABC program. They found that over the two years, 61.8-86.5% of the free-ranging population was sterilised and vaccinated. They predicted that by maintaining that level of ABC intervention, the dog population would decrease by 69% after 13-18 years and vaccination coverage would stabilise at over 70%. However, there has been no long-term data on any of the ABC programmes, making it difficult to assess the actual success of the ABC strategy.

In the Indian city of Bangalore which is the setting for this study, the ABC programme can reliably be considered unsuccessful. The most recent official free-ranging dog census, conducted in 2019, estimated the number at just over 0.3 million, compared to a human population of around 12 million (Worldwide Veterinary Service Centre, as reported by Sharma 2019). The census found that 46% of the free-ranging dogs in the city had not undergone the ABC programme. The report explained that the population had increased sharply from the 0.185 million dogs found in a 2013 census by the city’s administrative body – the Bruhat Bangalore Mahanagara Palike (BBMP).

Given the inadequacy of the sterilisation method, interventions premised on reducing the carrying capacity of the environment have been proposed. Totton et al. (2010) suggested that in addition to the ABC program, we must reduce the available food in the environment to assist in controlling the population. Baquero, Akamine, Amaku and Ferreira (2016) found that the latter is the most effective way to influence dog populations, compared to changing rates of abandonment, adoption, sterilisation and changing the carrying capacity of domestic dogs. This approach has been mildly echoed in several articles and directives which have proposed that the public can contribute to managing the population by not dumping waste in public (Herbert, Basha and Thangaraj 2012; KIMS 2007; Reese 2005) and by regulating slaughterhouses dumping waste (KIMS 2007). However, these directives operate on the assumption that refuse is the primary food source of free-ranging dogs. Meanwhile, other researchers propose that most of the dogs are in fact significantly dependent on ‘reference households’ or ‘reference individuals’ which actually support the bulk of the free-ranging population through direct feeding (Belsare and Gompper 2013; Cliquet et al. 2007; Morters et al. 2014; Reese 2005). Due to the sustained dependence of these dogs on specific houses, they are sometimes referred to as ‘owned’ dogs, despite being allowed to roam freely (Morters et al. 2014). The third proposed source is commercial areas which carry high quantities of organic material. Reese (2005) proposes that in North India there might be enough food sources from ‘food markets, slaughterhouses, temples and roadside restaurants’ to sustain the dogs. Yet, the nutritional dependence on the different food sources has not been quantified, because as Morters et al. (2014) have stated, quantifying the “uptake of environmental resources is generally not practicable” (p. 1097).

With a focus on further investigating the potential for this line of interventions in Bangalore, we pose the first research question of the study, *‘How can we predict free-ranging dog population sizes based on different types of food sources?’* We seek to identify the role played by garbage piles, households and commercial establishments, respectively.

Given the tight-interconnectedness between humans and free-ranging dogs, Matter and Daniels (2000) state that the most important determinant of dog population size is the attitude of the relevant communities. Similarly, any interventions that could be based on food sources would involve community-level changes. This requires a more in-depth understanding of community practises and opinions with respect to free-ranging dogs. However, the lack of data on strays extends to public opinion, which is primarily captured in the media, and therefore does not present a comprehensive view of attitudes, limiting the inputs to the framing of policy. While the media often reports opinions of upper-class residents, the perspectives of those who live closer to the dogs and are thereby affected more by them, is lacking. An exception is a paper by Herbert, Basha and Thangaraj (2012), who interviewed residents of slums – which see a disproportionate number of rabies cases from dog bites. They found that 66.5% of slum respondents saw free-ranging dogs as a problem, because they bark and create a nuisance (37.3%) and attack and bite people (29.2%). Most respondents felt that the duty of the dog population control is largely that of the governments. The authors suggested that this lack of a feeling of responsibility for the problem could mean that people are not aware of the part they can play by for example, avoiding dumping food waste (Herbert, Basha and Thangaraj 2012).

As Reese (2005) said, “the success of such control measures depends heavily on an understanding of the dog ecology and the nature of the dog-human bond in the locale under consideration.” (p. 58). Without an understanding of the anthropological context within which interventions might be suggested, possible ecological interventions are limited in their practicability. Therefore, in addition to this paper being an ecological study, in it we aim to build upon the knowledge of attitudes towards dogs, of residents from different socio-economic classes in Bangalore. To do so we pose the second research question *‘What are the differences in opinions on the free-ranging dog-human conflict across socio-economic groups of the Indian urban public? Further, how can these differences be understood?’*

## Methods - Ecology of free-ranging dogs

In order to answer the first research question regarding how different types of food sources can be used to predict the population of dogs, we collected raw data on the distribution of dogs and food sources, and ran a generalized linear analysis with multi-model inference. In this section, the process of data collection, organisation and analysis is described.

## Data collection

### Location

The data regarding dog populations and food sources was collected in northern Bangalore, India, spread over the localities of Jakkur, RK Hegde Nagar, Amruthahalli, Sahakar Nagar, Kodigehalli and Thindlu (Fig. S1). These were urban areas, characterised by differences in density and socio-economic profile, which were also captured to ensure a representative sampling.

### Unit areas

The spatial scale used in the study was 248*248 m^2^. This was deemed appropriate based on existing data concerning the maximum home range of free-ranging dogs in India, which one study found to be 544m in diameter (Pal, Ghosh and Roy 1998). Spatial resolution twice as fine was therefore considered adequate to capture the dynamics of dog distribution in relation to food sources.

QGIS version 3.4.3-1 (QGIS Development Team 2018) was used to create the 248*248 m^2^ grid map which was imported onto My Maps for use while surveying the dogs. 28 units, each with fairly homogenous socio-economic class profiles, were selected for survey.

### SE class classification

It is important to note that economic status is difficult to gauge accurately without income data. Moreover, most neighbourhoods experience gentrification at some point and are not entirely homogeneous. But because the sample areas were on the outskirts of the city, the character of a neighbourhood was relatively singular and easy to characterise. The classification was made as follows:

a. Lower socio-economic class: small houses with no cars or only company-owned taxis (the resident being the taxi driver) parked outside (*n* = 9).
b. Middle class: small independent houses without space for cars. If present, the cars were low-end models and parked out on the street (*n* = 6).
c. Upper class: large independent houses with one or more personal cars either in driveways or parked outside (*n* = 8).
d. Empty: areas where housing development was just beginning. Largely characterised by sparsely interspersed upper-class houses, makeshift houses of construction workers and empty plots of land (*n* = 5).

### Dog population survey

A free-ranging dog population survey was conducted using photographic capture-recapture sampling method. This technique involves repeated sampling of a population, wherein the individuals are recognised in each subsequent sampling not by artificial physical marks but by comparison with pictures from previous sightings. The surveys were conducted by moving a steady speed on a bicycle through all the streets in the unit area and recording sightings using the mobile application Geopaparazzi. The route followed in each unit was chosen based on the most efficient way to cover every single street and remained the same in each subsequent sampling. In this program for each sighting, an image of the dog was captured along with its location. At the end of each sampling session, a .kmz file was exported for processing on Google Earth Pro which displays the locations of each dog photographed in a sample, along with the images. Four surveys were conducted in each unit over the span of two consecutive days, in the morning between 6 30 and 9 30 and in the evening between 15 00 and 18 00. Only in the case of one unit, the number of samples was three rather than four, because the weather impeded one sampling session. These times allow for enough light for photography and have been found by other authors to produce the highest number of dog encounters (Tiwari et al. 2018). The surveys took place between 16 January and 22 January 2019 with 1,796 sightings recorded. Of these, 761 individual dogs were identified, with 324 being ‘uniques’ or dogs that were only sighted once in total. The full dataset is presented in Table S1.

### Food source mapping

The routes used in the dog survey were subsequently covered by bicycle again and Geopaparazzi was used to mark the location and nature of each food source present. For the garbage piles, in addition to the location, an image was taken for further analysis. The sources fell into six categories, namely (a) garbage, (b) small roadside shops, (c) bakeries, (d) butcher/fish shops, (e) restaurants, and (f) houses. The last food source – houses – was numbered using the base print of buildings in Google Maps. Since the surveys were conducted in predominantly residential areas and the number of commercial establishments were few in comparison, all the building base prints were counted without checking whether they represented a commercial or residential unit. Likewise, in the entire area surveyed, there were only four small apartment buildings counted, and each was counted as one house since the number of households within were unknown.

The shops, bakeries, meat/fish shops and restaurants were marked as potential food sources as it was hypothesised that dogs would either be able to find food in garbage bins or, in the case of shops and bakeries, might be fed scraps by the proprietors and consumers. The different types of establishments were analysed separately. The food source dataset is presented in full in Table S2.

## Data curation

### Data count

The dog survey data were imported as .kmz files onto Google Earth Pro and the images from each sampling session were used to map the dogs to each other to identify the number of unique individuals. Each dog was pinned at the first location it was seen and the number of repetitive sightings recorded. Typically, dogs were seen only within one unit, however if they were seen over multiple units, they were coded in the unit they were first sighted in. When the images were unclear, an informed judgement was made based on proximity and knowledge of the dogs. This data were reduced to a list of (a) total number of dogs spotted in a unit and (b) number of dogs that had only been spotted once, i.e., uniques (Table S1).

### Population estimates

The Application SuperDuplicates online tool was used to make probability estimates of the population based on the raw data (Chao, Colwell, Chiu and Townsend 2017). Tiwari et al. (2018) and Tiwari et al. (2019) found this to be the most efficient tool for estimating free-ranging populations in urban and rural India. This tool makes use of the number of uniques observed in an area to extrapolate the number of undetected individuals. While it was initially created to estimate species richness, in this study, each dog is considered as a unique species, as done in the review by Tiwari et al. (2019), to procure a reliable minimum population estimate. The application offers the use of ‘incidence data’ where one inputs the number of observed dogs, the number of uniques (dogs spotted only once) and the number of sampling units (4 for all except 1). The population estimate produced under chao.2est was used as the final population in the analysis (Table S1).

### Garbage selection and sizing

For a garbage pile to be considered a viable source, it was visually assessed for recent organic matter. This was to distinguish those that could be reliable food sources from garbage piles that were either inactive or created during construction and that contained primarily inorganic matter. Where garbage piles lined a road bordering a sampling unit, they were included in the unit’s analysis. However, garbage piles in neighbouring sampling units were discounted for the sake of clarity.

The viable garbage sources were recoded by size since there were large differences in that regard. Garbage piles that measured between 2*2m^2^ and 4*4m^2^ (typically found in empty plots between houses) were counted as 1 unit. Larger piles (typically found on the side of busy roads) were coded proportionally.

## Data analysis

### Differences between neighbourhoods

Two one-way ANOVAs were conducted to analyse the differences between mean population of free-ranging dogs as well as mean number of houses in neighbourhoods of different socio-economic classes. This was done with IBM SPSS Statistics 24 (IBM Corp. 2016) using the data collected.

### Estimating percentage of houses that feed strays

This was done using the following formula:

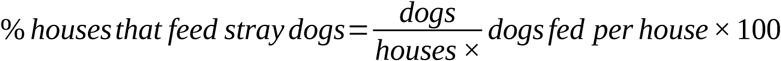

Number of dogs and number of houses were taken from the data collected per sampling unit, while the number of dogs fed per house was taken as 1.4. This number is based on figures produced by Morters et al. (2014). They found that in two villages in Johannesburg, South Africa as well as one in Bali, Indonesia, the average number of free-ranging dogs maintained by a single household was 1.3, and 1.7 in another village in Bali, producing an average of 1.4 overall. Since the average was sufficiently consistent over these widely different geographic and cultural contexts, we used this figure in our calculation.

### Generalized linear model analysis

Generalized linear models were constructed to analyse the weight of different food sources in predicting free-ranging dog populations. Since the response data had a count distribution which was overdispersed (μ_of population_ = 35.54, σ2_of population_ = 593.37), the GLMs were conducted in the form of negative binomial models with a log-link function. The analysis was performed in R 3.6.3 (R Core Development Team 2016). Model selection was carried out using the ‘MuMIn’ package (Bartoń 2016).

Prior to running the analysis, the predictor variables of interest were checked for multicollinearity. The variable ‘shops’ was removed from the dataset since the VIF exceeded the threshold value (VIF = 7.677). The adjusted dataset has low multicollinearity (VIF < 5). When interaction terms were added to this adjusted dataset, multicollinearity increased as expected. This was kept in mind while evaluating the model containing interaction terms.

### Model selection inference

Six alternative hypotheses were constructed to test potential relationships between various food sources and FRD populations. The hypotheses and the rationale behind them are detailed in Table I. Model selection and inference was applied by using the Akaike Information Criteria AIC_c,_ corrected for small sample size (n = 28). Akaike weights were used to evaluate the relative likelihood of the models. McFadden pseudo R^2^ is also presented to show the improvement of each model from null models.

**Table I.** Details of hypothesised GLM models included in analysis

| Hypothesis | Model | Predictors | Rationale |
| --- | --- | --- | --- |
| H1 | Direct sources | Bakeries, Houses | Sources at which people directly feed dogs |
| H2 | Indirect sources | Butchers, Restaurants, Garbage | Sources at which food (waste) can be found by dogs |
| H3 | Commercial sources | Bakeries, Butchers, Restaurants | Concentrated largely in commercial areas or streets |
| H4 | Non-commercial sources | Garbage, Houses | Not restricted to commercial areas |
| H5 | Direct sources and garbage | Bakeries, Houses, Garbage | Mixed group of direct and indirect sources to investigate whether the population depends on a combination to survive |
| H6 | Interaction between direct sources and garbage | Bakeries, Houses, Garbage, Houses*Garbage, Bakeries*Garbage | Investigate whether there is an interaction between direct sources ‘houses’ and ‘bakeries’ and indirect source ‘garbage’. Presence of significant interactions could indicate that garbage supplements direct sources in feeding some individuals of the population, in addition to the three sources independently supporting some individuals. |

In order to assess the relative importance of predictors in the best models, the Sum of Weights approach outlined in Burnham and Anderson (2002) was used. The AIC weights of the models where the predictor was present were summed up. In order to use this method, the number of models containing each predictor must be equal, so one model was added to this step to balance the set. The details are presented in the results.

## Methods – Differences in people’s attitudes

### Data collection

#### Participants

A sample of residents of northern Bangalore were surveyed about their attitudes towards free-ranging dogs. The participants were sampled from five socio-economic categories, namely Upper, Upper Middle, Middle, Lower Middle and Lower. Whereas in the ecological survey the categories were determined by the general socioeconomic character of the neighbourhood, here respondents were assessed individually. The lead author subjectively assigned the categorisation based on a combination of questions regarding the job of the main earner of the family, neighbourhood, ownership of a car or two-wheeler and type of house. A more reliable method would be to obtain the income of the household but due to the sensitive nature of the question, and the possibility that including it might have lowered the response rate or been answered untruthfully, the indirect method was used instead.

The upper-class respondents were mostly residents of a gated residential complex that the lead author grew up in. They were contacted by means of an email and answered the survey through Google Forms. The remaining respondents were recruited through convenience sampling in the street in various neighbourhoods that overlapped with the areas where the dog surveys were conducted. They were verbally asked the questions and their answers were recorded in the Google Forms document by the interviewer. One limitation of this methodological disparity is that it could account for longer and more deliberate answers being given by the first set of respondents, since they could spend more time on it while those in the street typically did not talk for long. Additionally, the respondents on the street were mostly spoken with in Kannada, which was then translated into English.

#### Survey

The ethnographic research was conducted by way of a survey which produced both quantitative and qualitative data (Survey questions available in Online Resource 1). The items on the survey related to the categories (a) Demographics, (b) Socio-economic class, (c) Opinion about free-ranging dogs and (d) Contact with (free-ranging) dogs. The quantitatively analysable questions that were used in the results were:

a. To what extent do you agree with the following statement: There is a stray dog menace that needs to be solved by the city administration. *1-5 Likert scale from ‘Disagree strongly to Agree strongly’.*
b. Do you think stray dogs should be removed from our cities? (*a) Yes (b) No (c) Maybe (d) Other.*
c. Do you feed stray dogs? *(a) Yes (b) No (c) No, but I support those who do.*

In addition, there were three opportunities for respondents to elaborate, which generated qualitative data that was used to explain the quantitative trends produced.

### Data curation

In order to analyse the results quantitatively, the responses to two questions needed to be recoded. First, the replies to the question ‘Do you think stray dogs should be removed from our cities? (a) *Yes (b) No (c) Maybe (d) Other’*, were recoded as Yes = 1, Maybe = 0.5, No = 0.

For respondents who did not give a clear ‘yes’ or ‘no’ answer, their answers were recoded into ‘yes’ or ‘no’ based on their longer opinion answers. Those with responses akin to, ‘we should not remove them all, but we should have fewer’ or ‘we should find a way to coexist’ or ‘only remove some – the dangerous ones’ – were recoded into ‘no’. Those whose other answers gave no further indication of how they felt, and one that essentially said, ‘they cannot be removed because of the impracticalities of current solutions’ was recoded as ‘maybe’. One that said dogs should be “removed from slums and areas where they are uncared for … should be tagged and only be kept in communities where individuals are willing to take responsibility to ensure they do not allow them to reproduce uncared (for) litters’’ was recoded as ‘yes’.

Second, for the question ‘Do you feed stray dogs? (a) *Yes* (b) *No* (c) *No, but I support those who do*’ the answers were recoded as ‘Yes = 1, No = 0’. The response ‘No but I support those who do’ was discounted since it is not informative of behaviour.

## Data analysis

### Quantitative analysis

The quantitative trends in responses were illustrated through a means plot and percentage-wise distributions of answers across socio-economic classes, to the three questions above. The figures were produced using the ‘ggplot2’ package (Wickham 2016) in R 3.6.3 (R Core Development Team 2016).

### Qualitative analysis

The qualitative analysis was performed on opinions expressed by survey respondents at three points in the survey. Views that were expressed repeatedly were highlighted and the number of times they were expressed by people from each class was assessed. These views were then categorised into three broad categories: ‘closeness’, ‘ethical reflection’, and ‘change and responsibility’. This semi-qualitative coding scheme was then drawn into an analysis that was used to explain the quantitative trends observed.

## Results – Ecology of free-ranging dogs

### Dog population statistics

The population of free-ranging dogs ranged from 192 to 1888 per square kilometre across the units sampled. The distribution of dogs was related to socio-economic class, with the lowest dog population found in the upper class neighbourhoods, with higher numbers in the middle, and highest in the lower-class areas (Table II). As can be expected, there is a corresponding increase in human population between the classes of neighbourhoods, as seen through the number of houses (Table III). There was a statistically significant difference across neighbourhoods with different socio-economic classes in the mean population of dogs (F = 9.52, p = 0.001) as well as the mean number of houses (F = 6.96, p = 0.005), as determined by conducting one-way ANOVAs (Table S6).

**Table II.** Mean population of dogs per class

| Neighbourhood Type | n | Mean # Dogs | Std. Deviation |
| --- | --- | --- | --- |
| Lower class | 9 | 57.4 | 27.42 |
| Middle class | 6 | 39.8 | 15.55 |
| Upper class | 8 | 17.0 | 3.07 |

**Table III.** Mean number of houses per neighbourhood type

| Neighbourhood Type | n | Mean # Houses | Std. Deviation |
| --- | --- | --- | --- |
| Lower class | 9 | 233 | 84.82 |
| Middle class | 6 | 176 | 43.48 |
| Upper class | 8 | 123 | 32.64 |

### Percentage of houses that feed free-ranging dogs

The average percentage of houses feeding free-ranging dogs was found to be greatest in the lower-class neighbourhoods, followed by the middle, and finally the upper-class areas (Figure 1). This difference between classes was found to be significant through a one-way ANOVA (F = 6.76, p = 0.006) (Table S6).

**Fig. 1.**
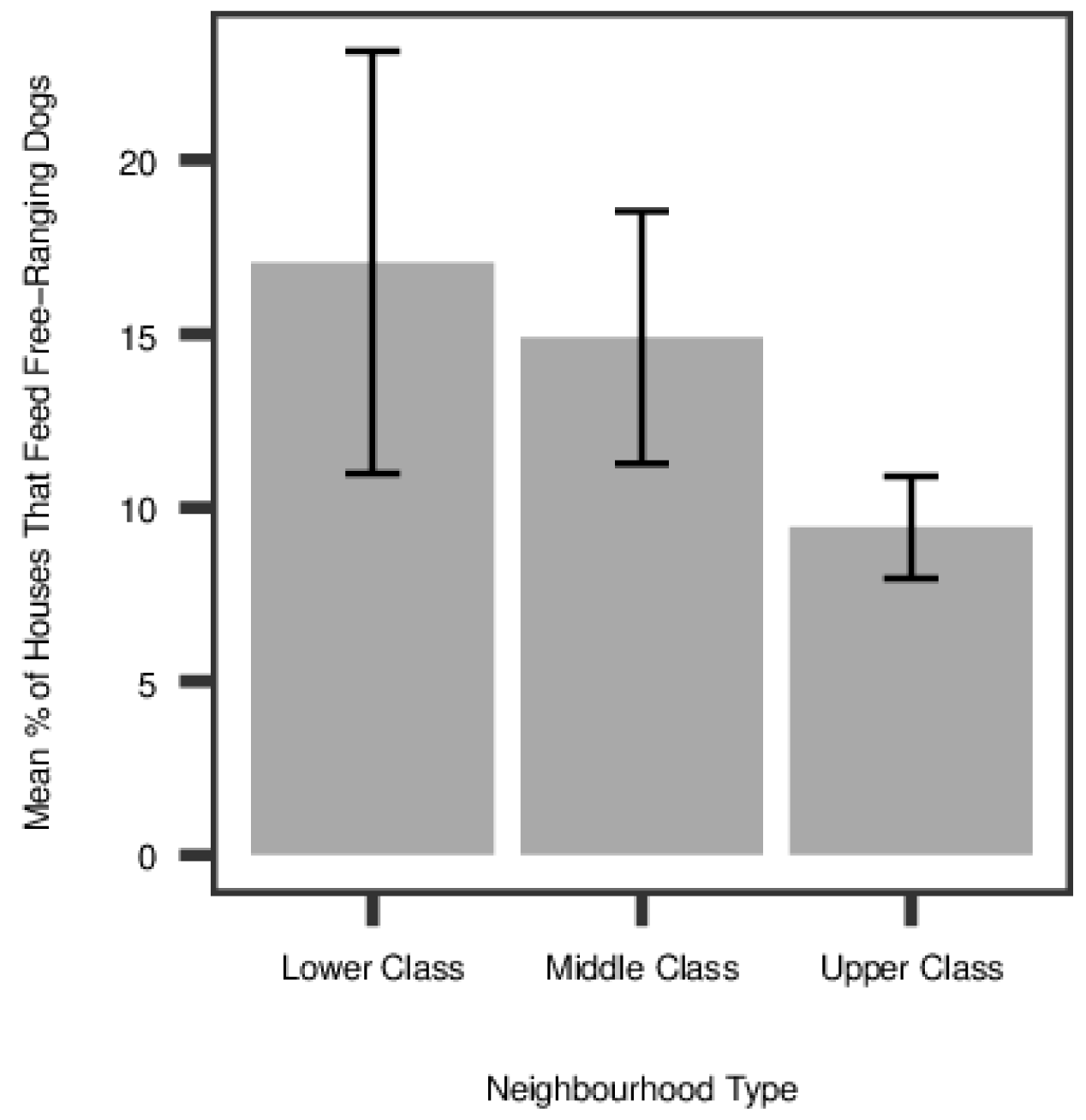
Means plot of % of houses that feed free-ranging dogs per neighbourhood type (*n* = 23)

### Generalized Linear Model (GLM) analysis with multi-model inference

The results of comparing six hypothesised GLMs by multi-model inference are presented below.

### Model selection and inference

The best models were ‘direct sources and garbage’ with AIC_c_ = 225.7, and ‘direct sources’ (AIC_c_ = 227.3), as shown in Table IV. Models ‘non-commercial sources’ (AIC_c_ = 228.9) and ‘interaction between direct sources and garbage’ (AIC_c_ = 229.7) were also ranked well (Table IV). However, due to the high multicollinearity in the interaction model(H_6_) and the little improvement over the ‘non-commercial sources’ model, the interaction terms were not considered important enough for inclusion. The remaining two models ‘indirect sources’ and ‘commercial sources’ had relatively little support in comparison (ΔAICc > 10). The goodness-of-fit measure shows that ‘direct sources and garbage’ (McFadden’s R^2^ = 0.214) has a strong predictive capability for free-ranging dog populations.

**Table IV.**
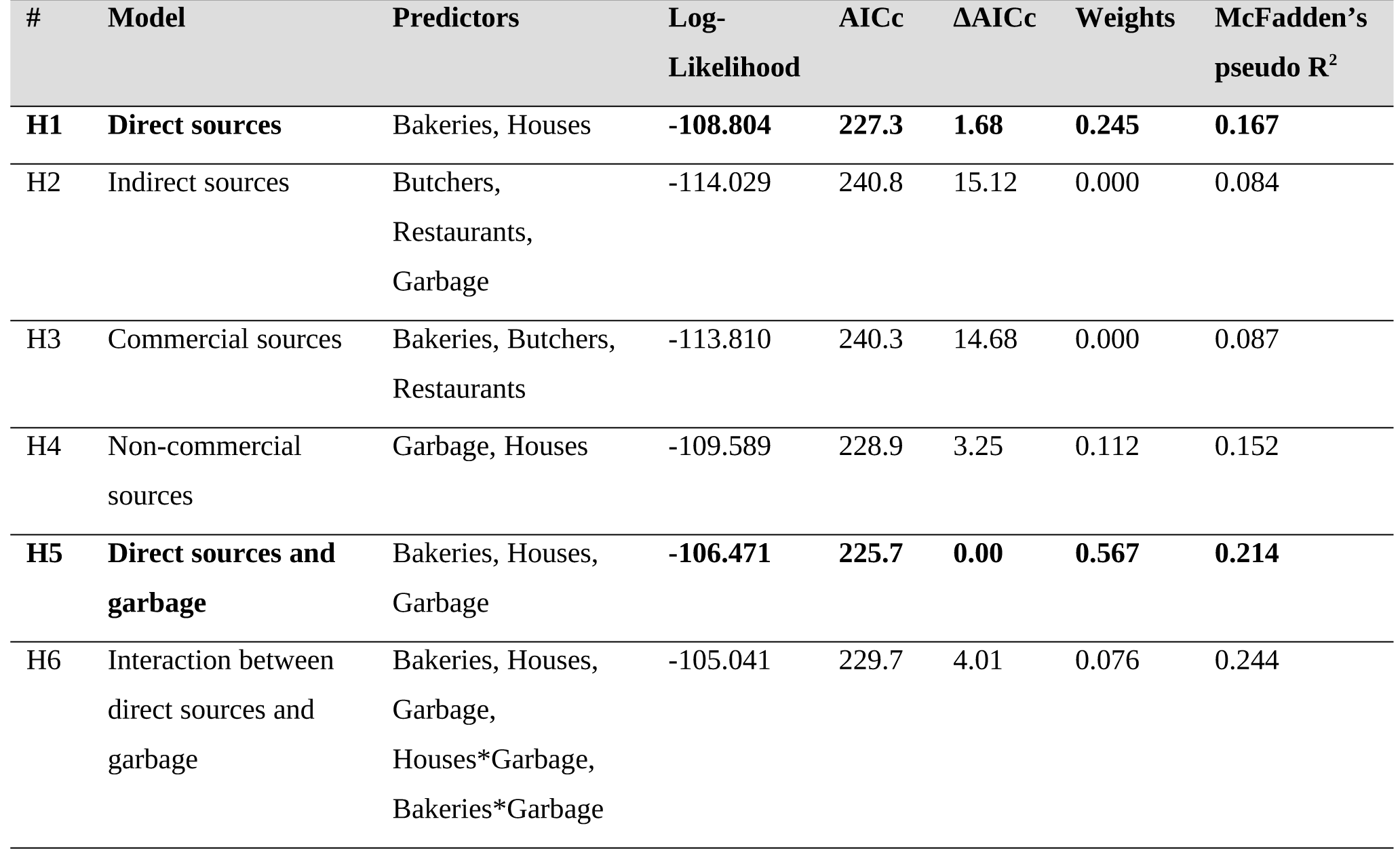
Results of model selection and multimodel inference as well as McFadden’s pseudo R^2^

### Parameter estimates for best models

The best model (H5,) contains a mixture of direct and indirect sources, and the second best model H1 comprises only direct sources. The parameter estimates and associated statistics for the best models are presented below (Table V and Table VI).

**Table V.**
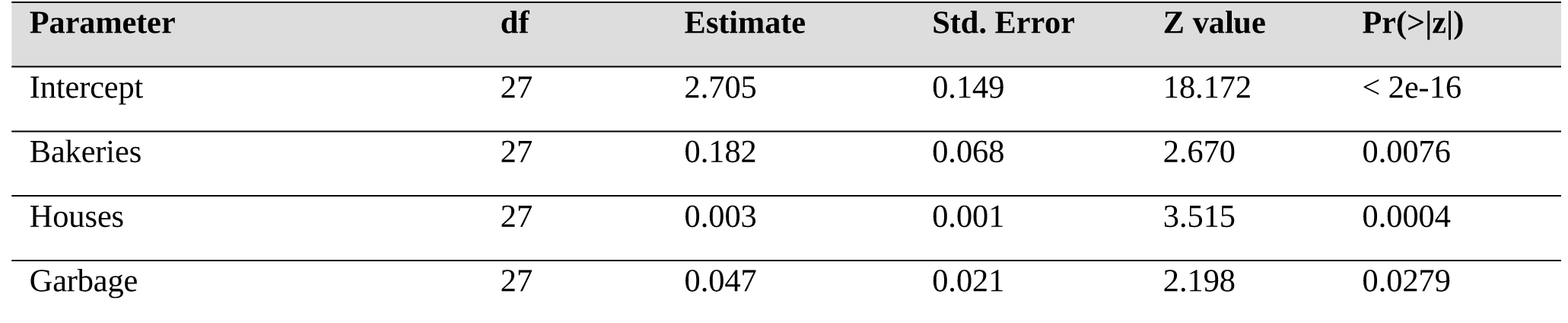
H5: ‘Direct sources and garbage’ model. Population = Bakeries + houses + garbage

**Table VI.** H1: ‘Direct sources’ model. Population = Bakeries + houses

| Parameter | df | Estimate | Std. Error | Z value | Pr(>\|z\|) |
| --- | --- | --- | --- | --- | --- |
| Intercept | 27 | 2.77 | 0.158 | 17.541 | <2e-16 |
| Bakeries | 27 | 0.163 | 0.076 | 2.150 | 0.0316 |
| Houses | 27 | 0.004 | 0.001 | 3.891 | 0.0001 |

From our best model we see that bakeries (p = 0.0076), houses (p = 0.0004) and garbage piles (p = 0.0279) are all significant predictors of free-ranging dog populations. In order to compare the relative weights of the three predictors, the best models were re-run, with the addition of a model containing bakeries and garbage to balance the set by ensuring that all the predictors were in an equal number of models (Table VII). The sum of weights for each predictor were then compared, showing that the houses were followed by bakeries and then garbage in importance (Table VIII).

**Table VII.** Models to determine relative importance of predictors

| # | Model | Predictors | Log-Likelihood | AICc | $\Delta AICc$ | Weights |
| --- | --- | --- | --- | --- | --- | --- |
| H5 | Direct sources and garbage | Bakeries, Houses, Garbage | -106.471 | 225.7 | 0.00 | 0.603 |
| H1 | Direct sources | Bakeries, Houses | -108.804 | 227.3 | 1.68 | 0.261 |
| H4 | Non-commercial sources | Garbage, Houses | -109.589 | 228.9 | 3.25 | 0.119 |
| - | Additional model with garbage and bakeries | Bakeries, Garbage | -111.517 | 232.8 | 7.10 | 0.017 |

**Table VIII.** Sum of Weights for predictors and relative importance

| Predictor | Sum of Weights | Importance rank |
| --- | --- | --- |
| Houses | 0.983 | I |
| Bakeries | 0.881 | II |
| Garbage | 0.739 | III |

## Results – Differences in people’s attitudes

A total of 100 individuals were surveyed for the ethnographic section of the paper, with 97 viable responses recorded. However, the sample size differs slightly per analysis because of missing responses on certain variables resulting in 84-97 responses per question. Of the 97 respondents there was almost an equal number of men (48) and women (49) (Table S3), and most were middle-aged, specifically 35-44 (17), 45-54 (33) and 55-64 (20) years of age (Table S4). Respondents were surveyed from across socio-economic classes, specifically Upper (40), Upper Middle (12), Middle (16), Lower Middle (9) and Lower (20) (Table S5).

The quantitative analysis of the survey responses produced certain trends in attitudes towards dogs. The upper class on the whole viewed free-ranging dogs more as a menace than the lower class, as seen in their response to the statement ‘There is a stray dog menace that needs to be solved by the city administration’ (Figure 2). Unexpectedly, the averages from the intermittent classes display a contrasting trend, as the upper middle classes viewed dogs less as a menace than middle and lower middle, which will be discussed further in the qualitative analysis.

**Fig. 2.**
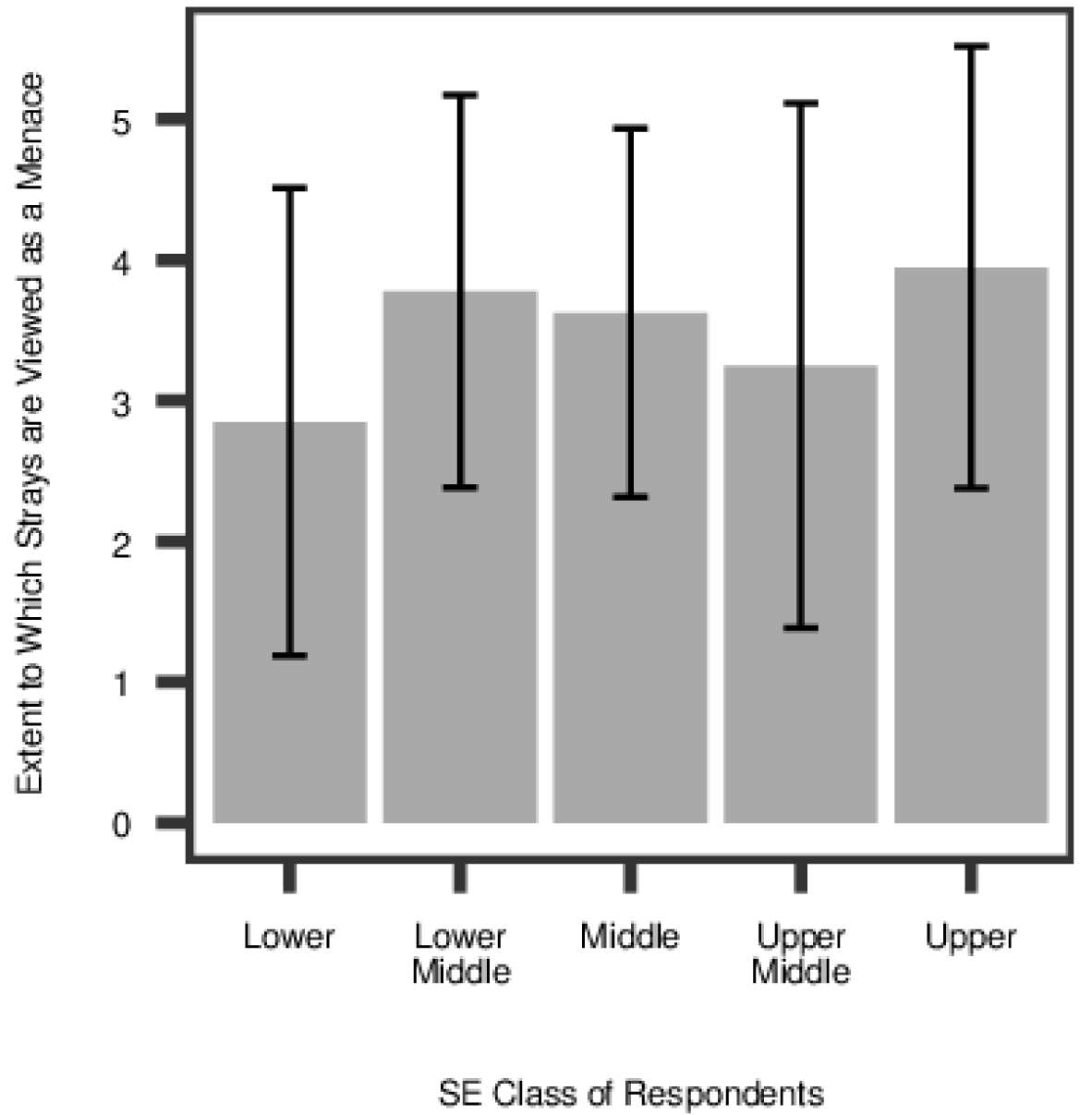
Means plot of extent to which respondents from different classes regard stray dogs as a menace to be solved by the city administration, on Likert scale from 1 to 5 (*n* = 95, *M* = 3.56, *SD* = 1.60). The standard deviation of each mean is represented as error bars

The second question ‘Do you think stray dogs should be removed from our cities’ was asked to assess the severity of opinions and how respondents viewed interventions (Figure 3). Over half of the individuals surveyed from each SE class responded ‘yes’ When contrasted with the responses to the previous question, this reveals a difference between how individuals from the lower and upper class perceive free-ranging dogs in general and their attitudes towards interventions, which is discussed further below.

**Fig. 3.**
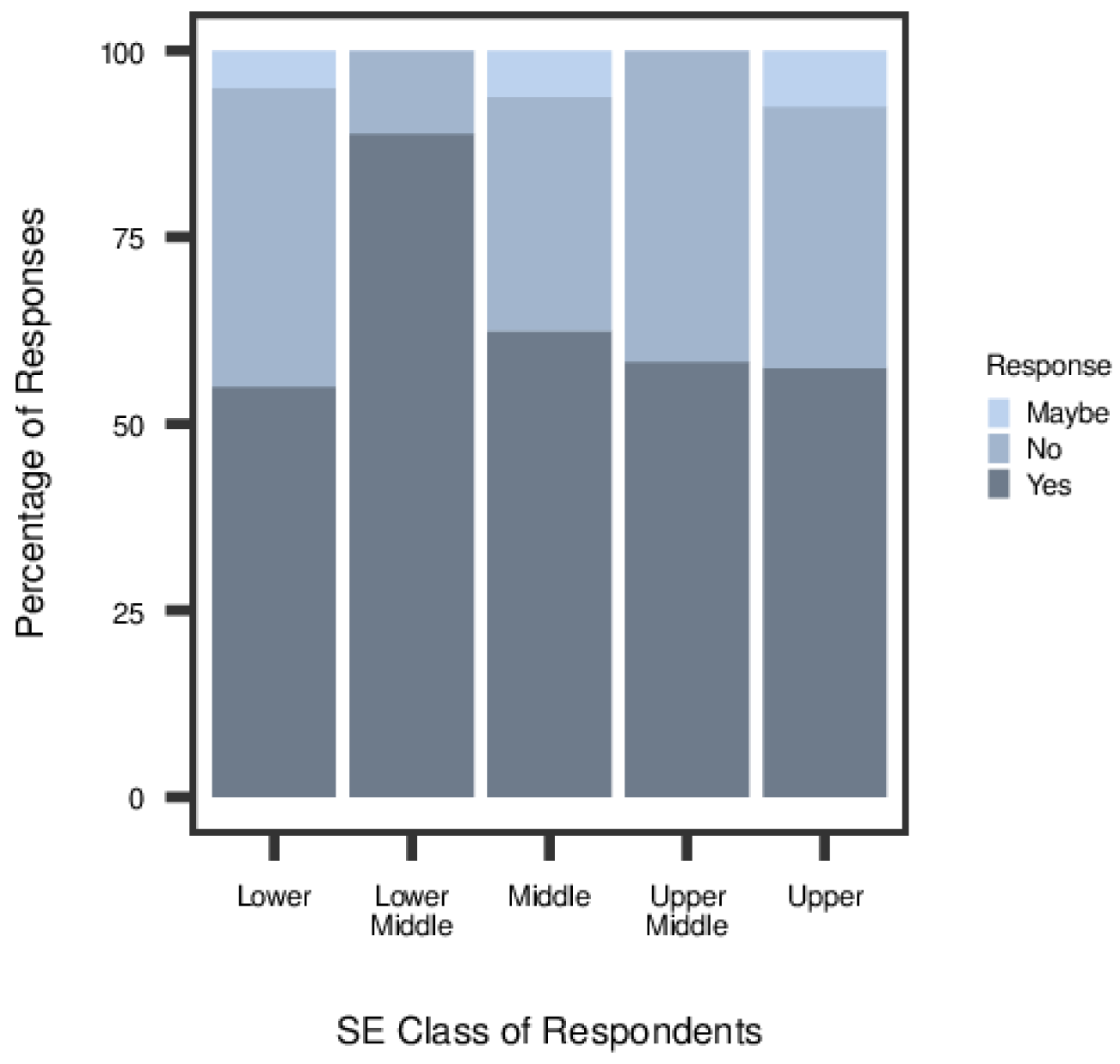
Percentage of respondents from each SE class who responded with ‘yes’, ‘maybe’ and ‘no’ to whether ‘free-ranging dogs should be removed from Indian cities’ (*n* = 97)

Finally, in order to investigate behavioural differences among the groups, we assessed what percentage of each class of respondents feeds free-ranging dogs. Figure 4 shows that high percentages of all the classes, except the upper-class, feed dogs. This difference could be explained by the fact that most of the upper-class respondents were residents of a gated colony, who would have to make the effort of travelling outside their compound to find dogs to feed.

**Fig. 4.**
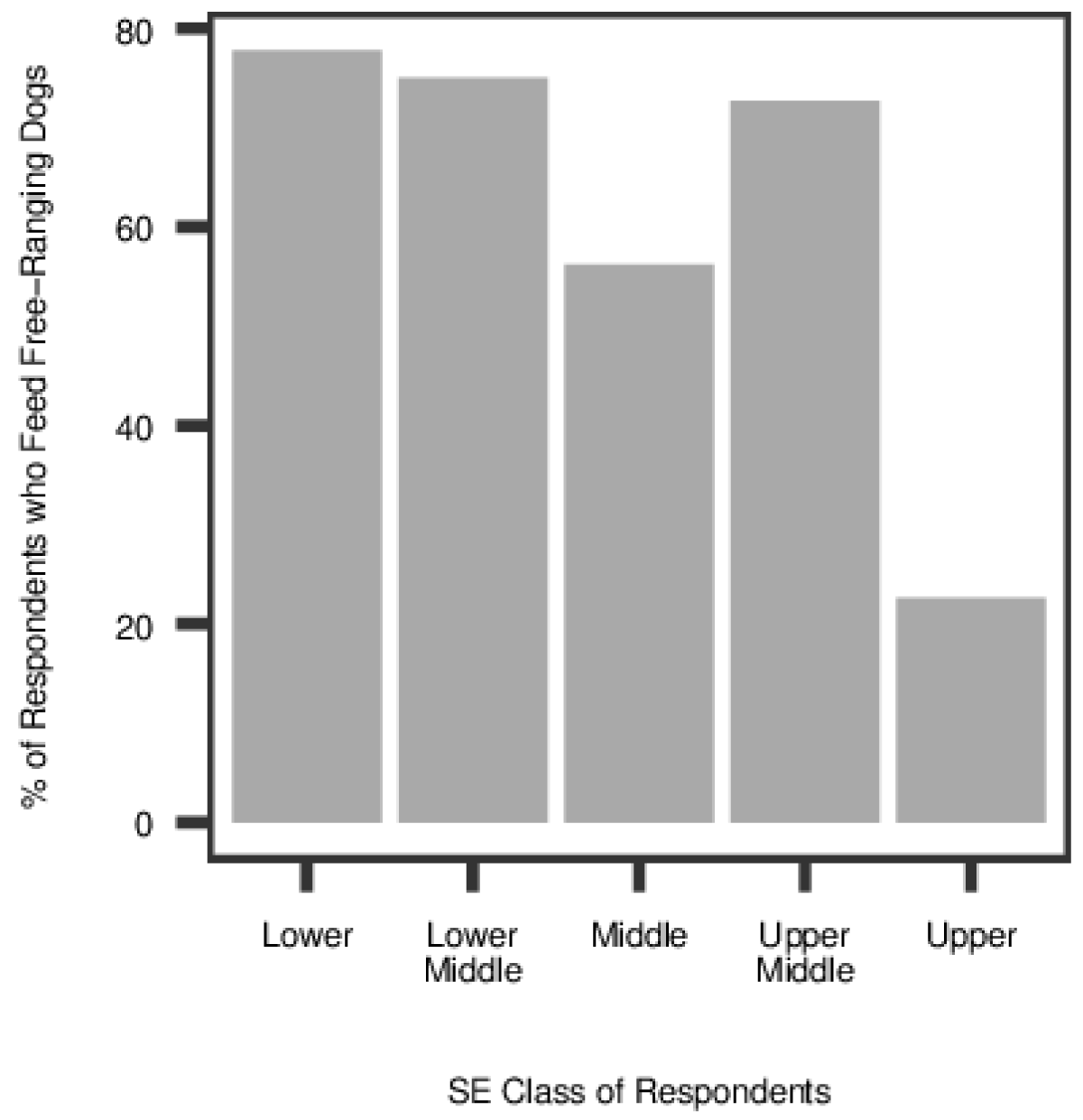
Percentage of respondents from different classes who report feeding free-ranging dogs (*n* = 84, *M* = 52.2, *SD* = 48)

### Qualitative analysis

The qualitative data obtained in the surveys is analysed in the following section and used to explain the results presented above. The findings are categorised into three themes that can be used to understand the differences in individuals’ opinions on free-ranging dogs. These themes are ‘closeness’, ‘ethical reflection’ and ‘change and responsibility’. The first of these describes differences that are associated with the physical ‘closeness’ between individuals and dogs, which is determined predominantly by where individuals live. The second and third themes explore differences that fall along socio-economic class lines.

### Closeness

Attitudes towards free-ranging dogs appeared to be strongly related to the physical closeness of the individual to the dogs. Closeness here is conceptualised as frequent daily contact between individuals and these dogs, as a result of living in close physical proximity to them. This was seen amongst individuals who lived in independent houses with gates that opened onto public streets, rather than those who lived in apartment buildings or gated colonies, who would not encounter free-ranging dogs in their immediate living environment. Importantly, this was not entirely determined by socio-economic class differences. In this sample, the upper-class respondents were largely from a gated colony, but most of the surveyed individuals from other classes lived in independent houses. In a broader sample, we would encounter a better mix of socioeconomic classes and closeness across respondents.

We explain the association with closeness as the less overall contact a person has with free-ranging dogs, the less positive contact they have with the dogs and thereby, the more negative their views towards dogs, and vice versa. For people with low closeness, the contact or information that they would have about these dogs would tend to come from a combination of transitional encounters with them – while travelling through streets they inhabit – and news reports. The former is often negative as the dogs chase individuals and vehicles that pass through their territory. The latter is also usually not just negative, but drastically negative, as it offers a one-sided narrative constructed around deaths and dog bites. Particularly in recent years, the reporting on dog attacks has intensified. This could cause such individuals to develop an abstract and skewed perception of the animals. They are conceptualised by this group as a “dangerous nuisance” that “needs to be controlled”.

On the contrary, people who live in independent houses that encounter dogs on their doorstep have more contact with them. They are seen as companions, in addition to noisemakers and car-chasers. In other words, they are not necessarily viewed as a problem but a daily feature of these individuals’ lives and therefore such individuals have more of a nuanced view and refer to specific dogs rather than the collective. As explained by one respondent: “There are some that don’t bother you and some that can get quite aggressive” (upper-class female, 22 y.o.). This would explain why such individuals often proposed that certain “dangerous” or “mad” individual dogs be removed, rather than all of them, a distinction that people from the gated colony do not make.

One important difference in views towards free-ranging dogs unexpectedly was not related to closeness but rather to socio-economic class lines. The upper and upper middle classes viewed the dogs more in terms of the danger they pose, citing dogs’ propensity to ‘attack’ far more than those from the lower classes, while the latter also appreciated their ‘protective’ behaviour: “If they are a menace elsewhere they should be removed but here there’s no problem. They act as our guards” (lower-middle female, 35-44 y.o.). This aspect is important in framing suggestions for policy because it reflects the vulnerability felt by the lower classes towards burglars that free-ranging dogs help to mitigate. This will be further discussed later in the paper.

### Ethical reflection

We see an interesting result in comparing Figures 2 and 3. The figures show that for many of the upper-class respondents there is a disconnect between regarding free-ranging dogs as a menace, and wanting them to be removed from the city. Relative to the lower class, on average the upper class was far more likely to consider the dogs to be a menace. However simultaneously, the percentage of respondents from the upper (58%) class who thought ‘stray dogs should be removed from our cities’ is very similar to that of the lower (55%) class. And of those from the upper class who did not think the dogs should be removed, nearly half (43%) did consider them a menace (Likert scale 4 or 5). This unwillingness to remove dogs can be explained in terms of how individuals think about animal welfare issues. It appears that persons from the higher socio-economic classes think about dogs more on ‘ethical’ and ‘humane’ grounds. Many are seen to think on the lines of “Any option we choose, we must avoid cruelty to every living being - including Dogs” (upper-class male, 65-74 y.o). On one extreme there is the ‘animal welfarist’ category of individuals who are against removal because they equate a human’s life to a dog’s life and believe that both species have a right to the planet. For instance, one respondent said, “We need to learn to live in harmony with the beings we share the planet with. If there are people out there who think they need to “exterminate” strays or another species for that matter, then their thinking is no different from that of religious terrorists who in the name of “cleansing” take innocent lives” (upper-class female, 18-24 y.o.). Another similarly said, “We need (to) handle this issue the way we are handling stray human beings. In fact [the dogs] pollute less, as far as our environment is concerned” (upper-class male, 65-74 y.o.). On the other end of the spectrum, there was one individual who said that mass culling humanely would be beneficial to both species since the dogs lead a hard life. In-between are respondents we call ‘transporters’ who suggested that free-ranging dogs should be removed from the cities and kept on a farm somewhere because they do not want to kill the dogs: “They should be moved elsewhere. I can’t recommend that they are killed, and the ABC isn’t working, the numbers only increase” (upper middle-class male, 65-74 y.o.). Then we identified the ‘impractical’ who suggested that the ABC programme is the humane option and simply needs to be implemented effectively: “Bbmp (Bangalore’s civic administrative body) should have sterilized dogs many years back, unfortunately they only did it partially. This is the scientific way of managing stray dogs” (upper-class male, 45-54 y.o.). Finally, there were the ‘practical sceptics’ who would wish the dogs to be removed but recognise that there is no viable strategy to do so at the moment: “They cannot be removed as the population is high and the solution inhuman. But there needs to be a comprehensive discussion on practices worldwide and solutions” (upper-class female, 45-54 y.o.).

Moving from the upper to the lower-class responses we saw almost no responses in the ‘animal welfarist’ category, showing that it is a type of thinking predominantly unique to the upper class. Additionally, the humane angle was decreasingly brought up, with a concurrent increase in ‘eradicator’ thinking – i.e., “If they’re left alone, they’ll bite everyone… It’s better if they’re taken at once” (lower middle-class female, 25-34 y.o.). There was an increase in respondents opting for immediate removal because “dogs bite” or even simply because “they are a nuisance.’’ However, at the same time, in the lower class, there were also a number of respondents who considered free-ranging dogs ‘harmless’ and even ‘helpful’ because they “keep burglars out”, a consideration largely missing among the upper-class respondents.

This consideration or lack thereof of ‘what is ethical’ can be seen clearly in the types of solutions proposed. Among the upper classes, there was a clear preference for ‘humane’ methods of removal such as the ABC program, while among the lower classes, for those who did want removal, there was a preference for quick methods such as mass culling. In Table IX we show the proportion of individuals who preferred the ABC program to those who preferred mass culling, amongst those who opted for ‘Yes they should be removed’. Those who suggested other methods or a mix of methods have not been included in this comparison.

**Table IX.**
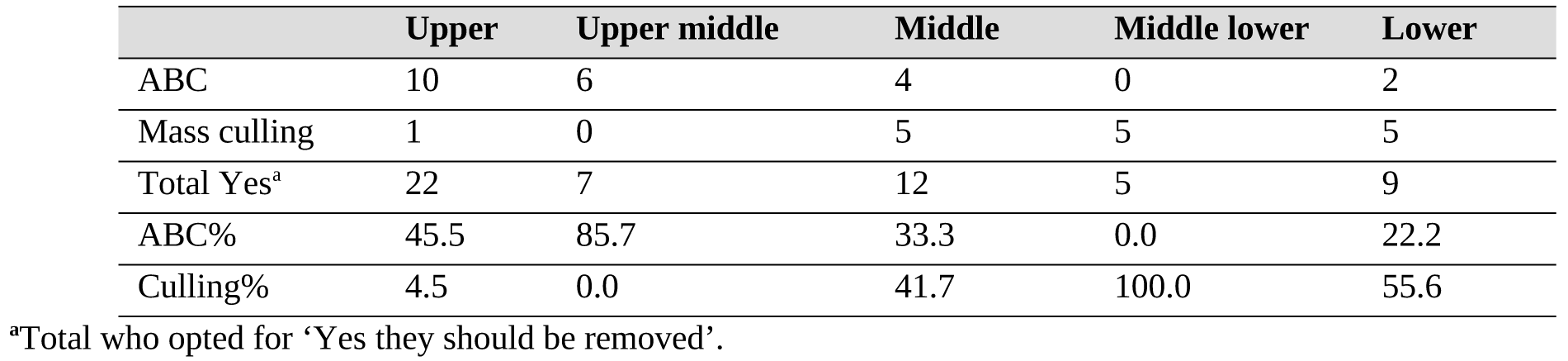
Preference for removal by ABC or Mass culling by class

|  | Upper | Upper middle | Middle | Middle lower | Lower |
| --- | --- | --- | --- | --- | --- |
| ABC | 10 | 6 | 4 | 0 | 2 |
| Mass culling | 1 | 0 | 5 | 5 | 5 |
| Total Yes<sup>a</sup> | 22 | 7 | 12 | 5 | 9 |
| ABC% | 45.5 | 85.7 | 33.3 | 0.0 | 22.2 |
| Culling% | 4.5 | 0.0 | 41.7 | 100.0 | 55.6 |

<sup>a</sup>Total who opted for ‘Yes they should be removed’.

### Change and responsibility

Besides the differences in ethical reflections, individuals from different socio-economic classes also tended to display variations in how they discussed the subject of interventions. For instance, respondents from the middle, lower middle and lower classes were less proclamatory in their proposed solutions than the upper classes: “If they can go, that will be good” (lower middle-class male, 65-74 y.o.). In the conversation it became clear to the interviewer that effective interventions were out of the former’s range of expectations. Therefore, solutions proposed were clearly spoken of in hypothetical terms or with scepticism, such as “If they could all be removed at once that would be great” (lower-class female, 55-64 y.o. & middle-class female, 45-54 y.o. & others). However, from context it was apparent that they were used to adjusting to conditions and had little faith that anything would truly change.

In contrast, the upper- and upper middle-class respondents proposed solutions with more self-assurance and vehemence in their beliefs, phrasing their responses along the lines of “something must be done”. It would appear that the upper classes felt less bound by the inefficiency of the administration in envisioning interventions. This could be because they are more insulated from the inefficiency on a daily basis, while the lower classes encounter it more, for instance in having to deal with a regular shortage of water. Even the ‘practical sceptics’ group among the upper class which proposed that there was no feasible solution yet often suggested a multi-pronged approach, maintaining that the dogs “need to be controlled”.

This also suggests a higher feeling of human responsibility for the situation amongst the upper classes who tended to view dogs as ‘for us to take care of’, while the lower classes might view them more as a part of the natural environment: “If you trouble them, they trouble you. If you leave them alone, they leave you alone. Why do anything to them?” (lower-class female, 45-54 y.o.). Further, there was also a variation in the assignment of responsibility. All the classes place responsibility for the intervention on the civil administration (BBMP), however in the upper, upper middle and middle classes, there was also mention of how citizens can contribute to the solution by not feeding or abandoning dogs. In contrast, there was a greater lack of perceived personal responsibility in dog population management from the lower classes, probably as a result of the closeness to dogs. This is particularly well illustrated by their feeding behaviour towards the dogs. Presented in Table X we can see that there was a disconnect between feeding free-ranging dogs and thinking that they should be removed, in the classes living closer to the animals. Although some respondents in the upper middle and middle classes displayed perceived personal responsibility in their responses, others displayed the same disconnect as the lower classes. Personal responsibility thus appears to be a feature of classes that is moderated by closeness to dogs.

**Table X.** Feeding behaviour*opinion on removing free-ranging dogs by class

| **Class** | **n** | **Don't**<br>**feed/Remove** | **Feed/**<br>**Remove** | **Feed/Don't**<br>**remove** | **Don't feed/don't**<br>**remove** |
| --- | --- | --- | --- | --- | --- |
| Upper | 26 | 19 | **0** | 5 | 2 |
| Upper Middle | 11 | 3 | **4** | 4 | 0 |
| Middle | 14 | 6 | **6** | 2 | 0 |
| Lower Middle | 6 | 2 | **2** | 2 | 0 |
| Lower | 19 | 3 | **6** | 9 | 1 |

This study aimed to understand differences in attitudes towards free-ranging dogs between socio-economic groups, and while class differences remain a prominent variable in understanding views, it appears that closeness to dogs was also an important factor. The responses of the upper middle class are a good way to identify what can be ascribed to the closeness variable and what is a feature of perspective based on rearing. Whereas closeness to dogs explains whether they are regarded more as a nuisance or menace, socio-economic divisions appear to explain the attitudes of people towards the ethics of specific interventions, the assertiveness with which they propose change and also whether they feel protected by the dogs. The perceived personal responsibility towards change was generally higher in the upper classes but tended to be lower among those from the same socio-economic group who live closer to dogs.

## Discussion

In this study, we have identified the food sources that substantially support the free-ranging dog population of North Bangalore. Additionally, we have extracted trends in attitudes of the residents from across socio-economic classes of the area. These results together allow us to propose an alternative approach to managing free-ranging dog populations in Bangalore by targeting the carrying capacity of the urban environment instead of the dog population directly.

We found that of the food sources surveyed, a mixture of direct sources and indirect sources significantly support the free-ranging dog population in Bangalore. Namely: houses, bakeries and garbage piles, in that order of relative importance. The foremost importance of houses confirms the findings by Butler and Bingham (2000), who likewise found that dog population density increases with an increase in human population. It also lends support to research that has found that free-ranging dogs in urban settings are supported primarily by referral households which engage in direct feeding of the animals (Morters et al. 2014). We found that the estimated percentage of households that feed these dogs is relatively small at 10% in upper-, 16% middle- and 18.3% in lower-class units, compared to 42 and 73% in two Indonesian villages (Morters et al. 2014). It is thus evident that a small proportion of houses can sustain large free-ranging dog populations.

The finding that garbage sources, although significant, are weaker predictors of populations goes against research by Butler and Bingham (2000) as well as Reese (2005) who suggested that in India, human waste food and faeces contribute highly to dog populations. We suggest that garbage serves as a secondary food source to household-maintained dogs (owned free-ranging dogs) and as the primary food source to the population that is not linked to specific households. Although the sampling methodology of this study did not allow for the quantification of the ratio of owned to unowned populations, Morters et al. (2014) found that only 1.1 to 1.5% of dogs in two study sites in Bali were entirely unowned, which suggests that the unowned population is likely to be small in this study area as well. During the data collection, it was also noticed that large garbage piles were not very common, contrary to popular opinion. Many neighbourhoods, particularly dense ones found in lower socio-economic areas did not have garbage piles in the narrow streets and the door-to-door garbage collection system appeared to be regular and well-used. Garbage piles were only seen on large roads, main shopping streets and in vacant lots present in middle- and upper-class neighbourhoods.

In terms of the attitudes towards free-ranging dogs and their control, we saw that, on the whole, the upper class respondents were far more likely than the lower class to consider them a menace that needs to be addressed. Additionally, we found three mechanisms that can be used to understand differences in opinion. Physical closeness to the dogs is a class-independent variable that can be linked with greater appreciation for the animals. On the other hand, socio-economic class does also play a role in explaining differences in how free-ranging dogs are viewed, in three ways: a. upper classes are likely to pay more attention to the ethical side of interventions than the lower classes, which were more likely to opt for quick methods of population removal; b. there is a greater belief in the plausibility of change, and the assertiveness with which interventions are suggested by the upper classes than the lower classes; c. the lower classes tend to feel less personal responsibility towards dog population management, in spite of (or perhaps because of) often living in closer proximity to them.

Nonetheless, although opinions differed widely across groups in the sample, there was a net recognition that free-ranging dogs pose an issue and require better management. Since there was no consensus on removing them from the city entirely, we propose reducing the carrying capacity of the environment and thereby the free-ranging dog population long-term. After all, even if the ABC programme were to be implemented more effectively, as long as the same amount of food sources are available, dogs from surrounding areas would enter the urban environment. Ultimately, reducing the population would allow the lower socio-economic classes to retain the protection of the dogs while reducing the ‘nuisance’ that they cause. In order to do so, we make two policy suggestions. First, in order to reduce the carrying capacity of the environment and thereby the long-term free-ranging dog population, we suggest regulating the feeding of dogs around bakeries and implementing proper waste management in public spaces. Second, we advise that by increasing awareness about the role that the household plays in sustaining a population, we can increase individuals’ feeling of personal responsibility in controlling the population, particularly amongst the lower classes, but generally in those who live close to the dogs. Since even the group that feels protected by the dogs often said that they would prefer fewer numbers, this awareness could also be useful in these neighbourhoods. While Herbert, Basha and Thangaraj (2012) suggested that households can be made aware of the role they can play by not dumping waste, we propose that more important is the awareness of the impact of feeding the dogs directly. An increased feeling of personal responsibility for owned free-ranging dogs might also contribute to sterilisation and immunisation coverage.

The value of the results and policy suggestions in this study can be verified by future interventional research that quantifies the impact on the population by reducing the environmental carrying capacity in the manner proposed here. Future planning would also be aided by gathering data on the ratio of owned to unowned free-ranging dogs in the Indian setting, as well as surveying the actual proportion of houses that feeds them. Additionally, since the study was conducted at the outskirts of the city, a survey area in the centre of the city is suggested to test the proposed theories. Likewise, the empirical evidence for the closeness hypothesis could be strengthened by surveying respondents from a mix of different housing types and socioeconomic groups.

## Conclusion

This paper looked at the impact of different food sources on dog populations as well as the dynamic between free-ranging dogs and humans from different socio-economic classes. We found that houses, bakeries and garbage piles were significant predictors of dog population sizes, and that there was a significant difference between dog populations in high and low socioeconomic neighbourhoods. Crucially, it was found that a small number of houses can support a large population of free-ranging dogs, while trash piles serve a secondary role in comparison. Across classes, opinions towards the animals differed quite widely but over half the respondents from each class felt free-ranging dogs should be removed from cities and the net opinion appeared to be that at the least, the free-ranging dog population needs to be controlled. Thus, we suggest that the city administration take steps to reduce the carrying capacity of the environment by regulating feeding around bakeries and improving waste management in public spaces. Creating awareness about the impact that household feeding has on the dog population could additionally help to control the population.

## Acknowledgements

The authors are grateful to Nachiket Kelkar and Abhijit Kumar for their helpful inputs to the paper.

## Declarations

### Compliance with ethical standards

Informed consent: Informed consent was obtained from all individual participants included in the study. Ethics approval for the dog population survey was not sought since the survey method was photographic capture-recapture from a distance, involving no direct interaction with the animals.

