## Supplementary material for "‘Stray Appetites’: A Socio-Ecological Analysis of Free-Ranging Dogs Living Alongside Human Communities in Bangalore, India"

### ESM

† Equal contributers

### Survey area

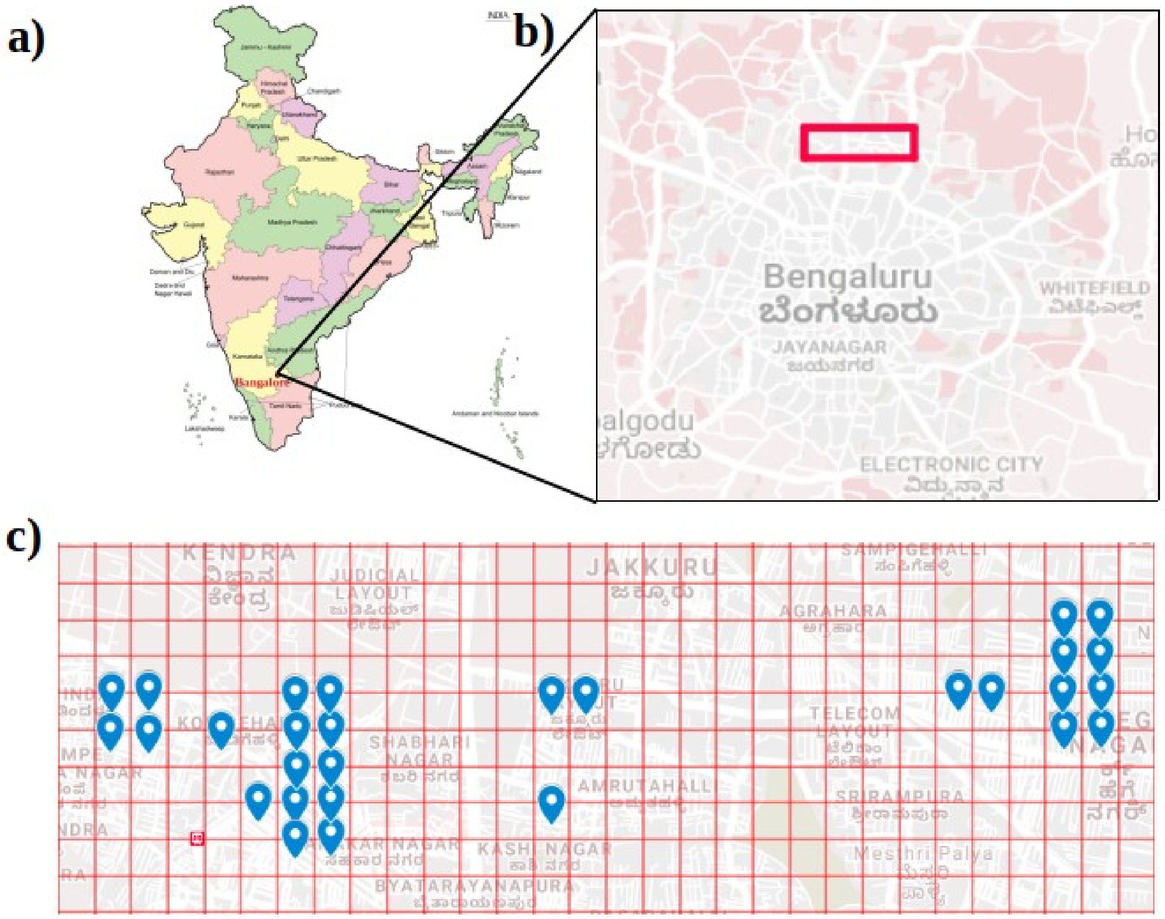


**Fig. S1** a. Position of Bangalore city on India map (Accessed from https://commons.wikimedia.org/wiki/File:India-map-en.png#/media/File:India-map-en.png). b. Area in North Bangalore, India where both surveys of free-ranging dog populations and qualitative surveys of residents were conducted for this study. c. Units of area 248*248m$^2$ where free-ranging dog populations were surveyed by photographic capture-recapture

### Ecology Data Tables

**Table S1.** Free-ranging dog population numbers in sampled areas

| Grid Reference | #Dogs sighted | #Uniques | #Samples | Population estimate (from Superduplicates) | se of population estimate |
|---|---|---|---|---|---|
| 1 | 14 | 9 | 4 | 24.16 | 8.63 |
| 2 | 21 | 9 | 4 | 27.04 | 4.28 |
| 3 | 38 | 19 | 4 | 52.92 | 7.35 |
| 4 | 65 | 27 | 4 | 82.54 | 7.86 |
| 5 | 32 | 16 | 4 | 44.54 | 6.69 |
| 6 | 9 | 7 | 4 | 21.68 | 11.45 |
| 7 | 15 | 5 | 4 | 17.7 | 2.66 |
| 8 | 14 | 7 | 4 | 19.46 | 4.19 |
| 9 | 11 | 6 | 3 | 15 | 4.81 |
| 10 | 12 | 4 | 4 | 14.16 | 2.55 |
| 11 | 11 | 5 | 4 | 14.54 | 4.1 |
| 12 | 18 | 4 | 4 | 19.65 | 2.01 |
| 13 | 12 | 5 | 4 | 15.26 | 3.88 |
| 14 | 26 | 12 | 4 | 34.63 | 5.36 |
| 15 | 14 | 9 | 4 | 24.12 | 10.92 |
| 16 | 9 | 5 | 4 | 13.48 | 5.56 |
| 17 | 23 | 11 | 4 | 31.22 | 5.27 |
| 18 | 32 | 12 | 4 | 39.13 | 4.06 |
| 19 | 11 | 3 | 4 | 12.41 | 1.81 |
| 20 | 16 | 3 | 4 | 17.12 | 1.46 |
| 21 | 28 | 12 | 4 | 36.06 | 4.79 |
| 22 | 37 | 10 | 4 | 41.67 | 3.28 |
| 23 | 50 | 21 | 4 | 63.76 | 5.72 |
| 24 | 95 | 37 | 4 | 117.69 | 8.11 |
| 25 | 42 | 19 | 4 | 55.36 | 6.4 |
| 26 | 45 | 22 | 4 | 61.84 | 7.95 |
| 27 | 26 | 9 | 4 | 31.01 | 3.47 |
| 28 | 35 | 16 | 4 | 46.4 | 5.86 |

**Table S2.** Population and food source data of areas sampled

| Grid # | Chao2.est | Garbage | Shops | Bakeries | Butchers | Restaurants | Houses | SE class |
|---|---|---|---|---|---|---|---|---|
| 1 | 24.16 | 2 | 1 | 0 | 0 | 0 | 113 | M |
| 2 | 27.04 | 1 | 1 | 0 | 0 | 0 | 176 | M |
| 3 | 52.92 | 1 | 2 | 1 | 0 | 0 | 182 | L |
| 4 | 82.54 | 2 | 7 | 2 | 1 | 1 | 229 | L |
| 5 | 44.54 | 6 | 4 | 0 | 0 | 0 | 143 | L |
| 6 | 21.68 | 2 | 1 | 0 | 0 | 0 | 144 | U |
| 7 | 17.7 | 1 | 2 | 0 | 0 | 0 | 112 | U |
| 8 | 19.46 | 0 | 3 | 1 | 0 | 0 | 126 | U |
| 9 | 15 | 0 | 0 | 0 | 0 | 0 | 111 | U |
| 10 | 14.16 | 2 | 1 | 0 | 0 | 3 | 125 | U |
| 11 | 14.54 | 3 | 0 | 1 | 0 | 1 | 88 | U |
| 12 | 19.65 | 1 | 1 | 0 | 0 | 0 | 189 | U |
| 13 | 15.26 | 3 | 2 | 0 | 0 | 0 | 75 | E |
| 14 | 34.63 | 0 | 1 | 0 | 0 | 0 | 24 | E |
| 15 | 24.12 | 3 | 1 | 0 | 0 | 0 | 25 | E |
| 16 | 13.48 | 4 | 0 | 1 | 0 | 0 | 89 | U |
| 17 | 31.22 | 0 | 1 | 1 | 0 | 0 | 180 | M |
| 18 | 39.13 | 1 | 3 | 0 | 0 | 0 | 157 | M |
| 19 | 12.41 | 1 | 0 | 0 | 0 | 0 | 67 | E |
| 20 | 17.12 | 3 | 0 | 0 | 0 | 0 | 46 | E |
| 21 | 36.06 | 0 | 0 | 2 | 1 | 3 | 135 | L |
| 22 | 41.67 | 0 | 6 | 1 | 1 | 5 | 287 | L |
| 23 | 63.76 | 2 | 16 | 1 | 1 | 3 | 389 | L |
| 24 | 117.69 | 6 | 21 | 5 | 15 | 7 | 315 | L |
| 25 | 55.36 | 9 | 3 | 0 | 0 | 1 | 184 | M |
| 26 | 61.84 | 11 | 5 | 1 | 0 | 2 | 247 | M |
| 27 | 31.01 | 5 | 8 | 0 | 1 | 0 | 244 | L |
| 28 | 46.4 | 11 | 5 | 0 | 0 | 3 | 177 | L |

### Descriptive Analysis of the Demographic Surveyed

**Table S3.** Sex descriptives of survey respondents

| Gender | Frequency | Percent | Valid Percent | Cumulative Percent |
|---|---|---|---|---|
| Female | 49 | 50.5 | 50.5 | 50.5 |
| Male | 48 | 49.5 | 49.5 | 100.0 |
| Total | 97 | 100.0 | 100.0 | |

**Table S4.** Age descriptives of survey respondents

| Age category | Frequency | Percent | Valid Percent | Cumulative Percent |
|---|---|---|---|---|
| 18-24 | 3 | 3.1 | 3.1 | 3.1 |
| 25-34 | 9 | 9.3 | 9.3 | 12.4 |
| 35-44 | 17 | 17.5 | 17.5 | 29.9 |
| 45-54 | 33 | 34.0 | 34.0 | 63.9 |
| 55-64 | 20 | 20.6 | 20.6 | 84.5 |
| 65-74 | 11 | 11.3 | 11.3 | 95.9 |
| 75-84 | 4 | 4.1 | 4.1 | 100.0 |
| Total | 97 | 100.0 | 100.0 | |

**Table S5.** Class descriptives of survey respondents

| Socio-economic class | Frequency | Percent | Valid Percent | Cumulative Percent |
|---|---|---|---|---|
| Lower | 20 | 20.6 | 20.6 | 20.6 |
| Lower middle | 9 | 9.3 | 9.3 | 29.9 |
| Middle | 16 | 16.5 | 16.5 | 46.4 |
| Upper middle | 12 | 12.4 | 12.4 | 58.8 |
| Upper | 40 | 41.2 | 41.2 | 100.0 |
| Total | 97 | 100.0 | 100.0 | |

### Survey questions

Demographic Variables:

1. Sex
2. Age

Economic background:

3. What is the job of the main earner of your family?
4. What is your type of residence?
5. Is your house owned or rented?
6. Which locality do you live in?

Opinion on dogs:

7. To what extent do you agree with the following statement: There is a stray dog menace that needs to be solved by the city administration. *1-5 likert scale from 'Disagree strongly to Agree strongly'.*
8. To what extent do you agree with the following statement: Stray dogs are not a problem and should be left alone by the city administration. *1-5 likert scale from 'Disagree strongly to Agree strongly'.*
9. Please **elaborate** on your opinion about stray dogs
10. Do you think stray dogs should be removed from our cities? a. *Yes* b. *No* c. *Maybe* d. *Other.*
11. Please **elaborate**.
12. If yes, how do you think they should be removed? a. *ABC program* (gradual sterilisation over years) b. *Mass culling* c. ***Other***.

Contact with dogs:

13. Do you feed stray dogs? a. *Yes* b. *No* c. *No, but I support those who do.*
14. Have you now or previously had pet dogs at home? a. *Yes* b. *No.*
15. For the most part, how do you get to work/school? a. *Walk* b. *Cycle* c. *Motorised two-wheeler* d. *Car* e. *Bus* f. *Train* g. *Other.*
16. Have you or has someone close to you (family or friend) ever been: a. *Bitten* b. *Chased repeatedly,* by a stray dog?

#### Mean differences between classes

**Table S6.** ANOVA of mean differences between socioeconomic units on % households that feed stray dogs, dog population and number of houses

| | | Sum of Squares | df | Mean Square | F | Sig. |
|---|---|---|---|---|---|---|
| % of Households That Feed Stray Dogs | Between Groups | 292.513 | 2 | 146.257 | 6.757 | .006 |
| | Within Groups | 432.888 | 20 | 21.644 | | |
| | Total | 725.402 | 22 | | | |
| Dogs | Between Groups | 6935.406 | 2 | 3467.703 | 9.516 | .001 |
| | Within Groups | 7288.392 | 20 | 364.420 | | |
| | Total | 14223.798 | 22 | | | |
| Houses | Between Groups | 51786.771 | 2 | 25893.385 | 6.955 | .005 |
| | Within Groups | 74459.056 | 20 | 3722.953 | | |
| | Total | 126245.826 | 22 | | | |
